# In search for the most optimal EEG method: A practical evaluation of a water-based electrode EEG system

**DOI:** 10.1101/2021.04.28.441825

**Authors:** Marta Topor, Bertram Opitz, Philip J. A. Dean

## Abstract

The study assessed a mobile electroencephalography (EEG) system with water-based electrodes for its applicability in cognitive and behavioural neuroscience. It was compared to a standard gel-based wired system. EEG was recorded on two occasions (first with gel-based, then water-based system) as participants completed the flanker task. Technical and practical considerations for the application of the water-based system are reported based on participant and experimenter experiences. Empirical comparisons focused on EEG data noise levels, frequency power across four bands (theta, alpha, low beta and high beta) and event-related components (P300 and ERN). The water-based system registered more noise compared to the gel-based system which resulted in increased loss of data during artefact rejection. Signal to noise ratio was significantly lower for the water-based system in the parietal channels which impacted the observed parietal beta power. It also led to a shift in topography of the maximal P300 activity from parietal to frontal regions. It is also evident, that the water-based system may be prone to slow drift noise which may affect the reliability and consistency of low frequency band analyses. Practical considerations for the use of water-based electrode EEG systems are provided.

## 1. Introduction

Brain activity measured using electroencephalography (EEG) allows for a close investigation of electrical signals in the frequency and time domains. It is a common neuroscience method used in psychological, behavioural, cognitive, and clinical research due to its affordability and ease of use. The standard state-of-the-art EEG equipment involves a swim-cap like device with inserted electrodes. The connection between the scalp and the electrode is normally bridged with electrolyte gel. The signal is recorded through electrode wires connected to a computer. Technological advancements aim to improve the useability of EEG systems through new electrode types and wireless EEG signal recording.

The main disadvantage of the gold standard gel-based electrodes is the time-consuming preparation process including skin abrasion, gel application and impedance checks. The setup time depends on the number of included electrodes and researcher experience but typically varies between the average of 30 to 70 minutes (Kam et al., 2019; Oliveira et al., 2016). After the recording, participants have to wash the gel out of their hair and the electrodes require cleaning. Skin abrasion and lengthy preparation may be problematic for participants with sensory sensitivities, attention difficulties and restlessness which are often observed in young children and individuals with neurodevelopmental conditions such as autism spectrum disorder (ASD) or attention deficit hyperactivity disorder (ADHD). The use of the systems is also limited to trained researchers or clinicians, but researchers often look for systems that can be applied independently by participants or patients at home (Hinrichs et al., 2020; Jochumsen et al., 2020; Radüntz, 2018).

Alternatively, systems that do not require skin abrasion such as EEG nets might be preferred (DiStefano et al., 2019; Pierce et al., 2021) though they generally use saline or gel solutions and cannot be used by participants independently. In the last few years, new systems with dry electrodes emerged. They require no skin abrasion, no gel/saline solutions and could potentially be used by participants independently without the presence of a trained researcher (Hinrichs et al., 2020; Kam et al., 2019; Pinegger et al., 2016). However, the data recorded with these systems have been reported to contain a higher number of artefacts (Hinrichs et al., 2020; Oliveira et al., 2016), higher pre-stimulus noise levels (Hinrichs et al., 2020; Mathewson et al., 2017) and lower signal-to-noise ratio (Radüntz, 2018) than standard gel-based electrode recordings. This is likely caused by the lack of an electrolyte substance that could bridge the scalp-electrode connection and keep the electrodes close to the skin throughout the recording (Mathewson et al., 2017; Pinegger et al., 2016). In addition, two studies reported lower participant comfort ratings for dry compared to wet EEG electrode systems due to the pressure from electrodes’ metal pins (Kam et al., 2019; Oliveira et al., 2016). These issues may deem dry-electrode systems unsuitable for many research designs.

Water-based electrodes are a promising development which could potentially improve on the gel-based systems’ disadvantages and mitigate the issues observed in dry-electrode recordings. They consist of plastic casings and paper or felt inserts soaked in tap water. Compared to gel-based electrodes, there is no need for skin preparation or washing hair and the preparation procedure is relatively easier and less time consuming. In contrast to dry electrodes, the scalp-electrode connection is supported with water which may help to sustain high quality signal. No metal parts of the electrodes come into direct contact with the skin thus potentially improving participant comfort as well. So far, the quality of EEG recordings using water-based electrodes has been evaluated in the context of brain computer interface (BCI) designs and the results are promising. Noise levels during a short circuit recording were the lowest in a water-based compared to gel-based and dry electrode systems (Pinegger et al., 2016) and the signal to noise ratio has been reported to be comparable between water- and gel- based systems (Jochumsen et al., 2020). In addition, participant satisfaction was the highest for water- compared to gel-based and dry systems (Pinegger et al., 2016). Moreover, the available water-based EEG systems allow for mobile wireless recordings of the EEG signals. This creates an opportunity to obtain EEG recordings in a wider range of contexts outside of the lab including everyday life situations, at home recordings, motor and sports research (Hinrichs et al., 2020; Oliveira et al., 2016; Radüntz, 2018). Taken together, water-based electrode EEG systems may seem very attractive for a wide range of research designs in neuroscience (examples of recent studies: Hazarika and Dasgupta, 2018; Raj et al., 2020).

To our knowledge, there are currently no empirical studies investigating the suitability of the new mobile water-based EEG electrode systems for application in cognitive and behavioural neuroscience research. We aimed to fill this gap by evaluating the quality of signal obtained with a water-based electrode system and investigating whether it may impact time-frequency and event-related potential (ERP) analyses. We also aimed to understand potential drawbacks and best methodological practices for the use of such systems. We applied evaluation methods and suggested benchmarking comparisons previously used for dry electrode systems evaluations (Hinrichs et al., 2020; Kam et al., 2019; Mathewson et al., 2017; Oliveira et al., 2016; Radüntz, 2018). Practical advice is provided alongside the obtained results for researchers who might want to consider using water-based electrode EEG systems in the future.

## 2. Method

### 2.1. Participants

The study consisted of two phases. Phase one was part of a procedure for an earlier study which used a gel-based EEG system (Topor et al., 2021). It included 46 participants in the pilot and the final study stage. Recruitment was facilitated through the University of Surrey’s research volunteer system and through word of mouth. All participants were given an opportunity to win one of two £50 prize vouchers and psychology students received lab tokens required as part of their course. Participants were excluded for diagnoses of psychiatric, neurological, or neurodevelopmental disorders.

In phase two, the same participants were contacted and invited to participate again. They were contacted one by one, in no particular order, and recruitment stopped when the 10^th^ participant agreed to complete the study. Sample size was determined based on previous evaluations of EEG systems which included eight to nine participants (Mathewson et al., 2017; Oliveira et al., 2016; Pinegger et al., 2016), and detected significant differences between the devices used. We also used a within-subject design which is advantageous for the preservation of power in studies with small sample sizes (Charness et al., 2012).

The final sample consisted of 10 participants who completed both phase one and two. We attempted to ensure a gender balance within the sample, and thus recruited five males and five females. Mean age at phase one was 26.5 years old, range of 22-38. The time between participation at phase one and two ranged from seven to twelve months. In the final sample, there were no individuals who won the £50 prize in phase one and no additional incentives were offered at phase two. No additional demographic or health checks were carried out at phase two. All participants provided written informed consent. The study complied with ethical regulations at the University of Surrey, approval ID 428470-428461-48044088.

### 2.2. Materials and Equipment

#### 2.2.1. Gel-based EEG system

The EEG recordings at phase one were acquired using the EasyCap gel-based setup (EasyCap system kit, Brain Products, n.d.; from now on referred to as EasyCap) with 32 Ag/AgCl sintered electrodes in a 10/20 system (Fp1, Fp2, Fz, F3, F4, F7, F8, FC1, FC2, FC5, FC6, Cz, C3, C4, T7, T8, CP1, CP2, CP5, CP6, TP9, TP10, Pz, P3, P4, P7, P8, POz, O1 and O2). The ground electrode was located within the cap at position AFz. The electrooculographic signal was recorded from the left side (vertical, VEOG) and above (horizontal, HEOG) the left eye using additional electrodes outside of the cap. The reference electrodes were also external to the cap, located on the mastoids and recorded implicitly (i.e., not as separate channels). Data were recorded in DC mode using Brain Vision Recorder V1.2 (Brain Products, 2012) at 500Hz with amplifier input impedance at 10GΩ and electrode impedance below 5kΩ. A high cut-off online filter was implemented at 250 Hz.

During the equipment preparation, each participant’s head circumference was measured with tape to select the right EEG cap size (52, 54, 56 or 58 cm available). The electrodes remained fitted within the caps between different recording sessions. Once the caps were placed on the head, the position of the electrodes was adjusted. The external electrooculographic and reference electrodes were placed on the skin using electrode stickers. Using a cotton bud, participant hair was moved from under the electrodes. We also applied an alcohol solution on the skin and the scalp directly through the hole in the electrodes (see figure 1 for an illustration). This was followed by the application of the electrolyte gel directly at electrode locations. In the case of noisy channels, it was possible to improve the signal quality by reapplying the gel and securing electrodes closer to the scalp. The preparation of each participant for recording lasted from 30 minutes to one hour.

**Figure 1.**
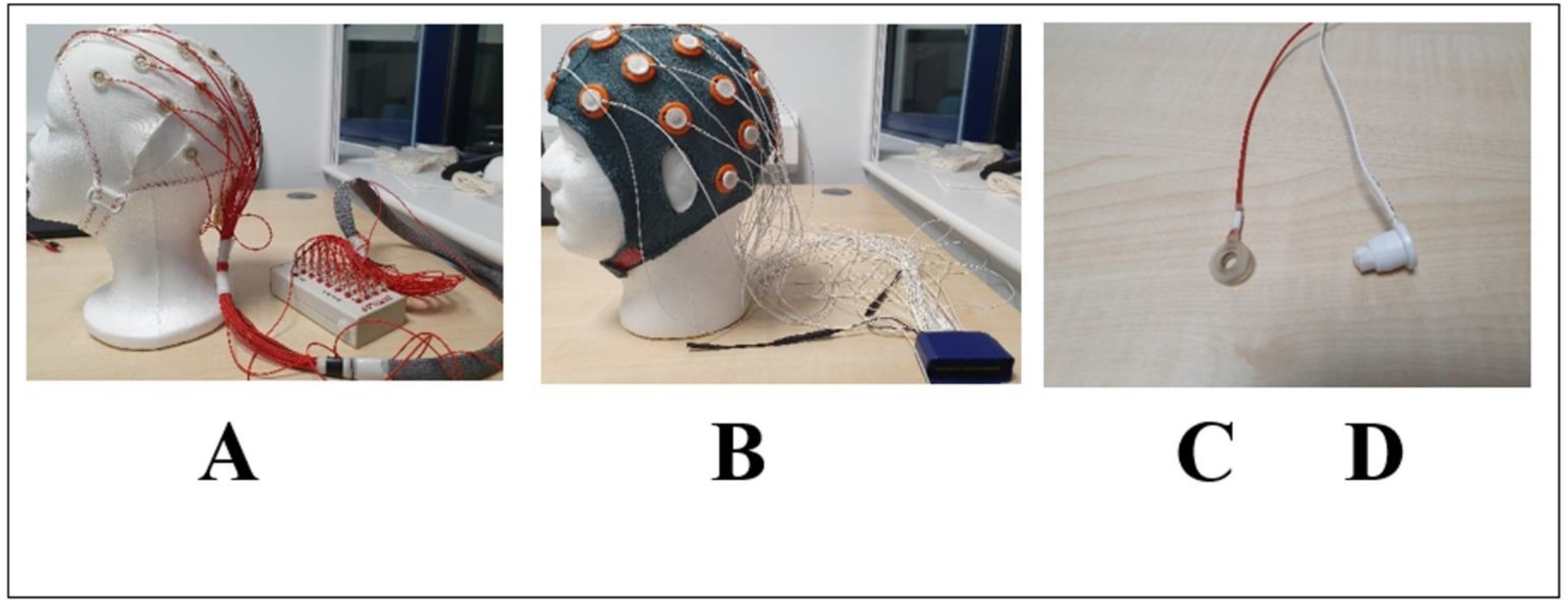
Two types of EEG recording setups and electrodes used in the current study. **A.** EasyCap setup. **B.** Mobita setup. **C**. A gel-based EasyCap electrode. **D**. A water-based Mobita electrode.

The electrode cables were gathered at the back of the participant’s head in a tight net and plugged in an input box connected to Brainamp MR Plus amplifier (Brain Products, n.d.). The EEG signal was recorded directly to a laptop that the amplifier was connected to using an USB adapter. The same USB adapter received the stimulus-response digital event markers from the stimulus computer via its parallel port. This wired setup enabled each stimulus and response category to have a unique signal at the parallel port translated to a unique marker value within Brain Vision Recorder alongside the EEG recording. All recordings were performed in an electrically shielded room.

#### 2.2.2. Water-based EEG System

During phase two, EEG data were acquired using the Mobita water-based setup (Mobita – W – 32 EEG, Biopac Systems, Inc., n.d.; from now on referred to as Mobita) with 32 electrodes in a Mobita-32EEG-CAP-A ConfiCap (Biopac Systems, Inc., n.d.). Similar to the EasyCap cap, it had fixed electrode positions in the 10/20 system, though included Fpz and Oz channels instead of TP9 and TP10. In contrast to EasyCap, the ground electrode was not located within the cap but secured with a sticker in the middle of the forehead. It was not possible to add more external electrodes so the electrooculographic signal was extracted from Fp1 for VEOG and F7 for HEOG. The reference electrodes were located on the mastoids within the cap and recorded as separate channels. Recordings were obtained in DC mode at 1000 Hz using the Acqknowledge software V 5.0.3 (Biopac Systems, Inc., 2018). The Mobita system does not allow for the measurement of electrode impedance. It has been argued that electrode impedance may have little influence over data quality if amplifier input impedance is high (Ferree et al., 2001). However, Mobita’s input impedance was comparable to that of EasyCap amplifier (10GΩ) so we aimed to monitor possible noise interference for consistency between the two systems. Therefore, live spectral power was visually inspected for each electrode to detect noisy spikes at 50Hz. Online filters were not applied.

Equipment setup included manual preparation of Mobita electrodes before participant arrival. Small pieces of absorbent paper (supplied by Biopac Systems, Inc.) were rolled and inserted into the plastic electrode casings. Electrodes were then placed in a jug of tap water. One adjustable cap size (medium: 54-58cm) with empty holes (grommets) for the electrodes was fitted and adjusted for correct positioning for all participants. Though skin preparation is not suggested in the Mobita instruction manual (Biopac Systems, Inc., 2019), we decided to apply the same alcohol solution used in the EasyCap recording in the areas with exposed skin (forehead and mastoids) to remove the natural oiliness which could prevent good conductance for the water-based electrodes. However, alcohol makes the skin dry which could also reduce the skin to electrode connectivity in water-based systems (Biopac Systems, Inc., 2016), so we did not apply it anywhere else. Before inserting the electrodes, participant hair was moved with a cotton bud to expose the scalp within the empty grommets. If a noisy spike was observed at 50Hz whilst checking the channels’ live spectral power, the electrode was removed, the hair was moved again to expose the scalp more and improve the electrode to scalp contact. In one case, due to noise across a number of electrodes, we tied a bandage around the participant’s head to keep the electrodes close to the scalp and prevent them from being dislocated by hair movement. Lastly, the Mobita amplifier (Mobita-W-32EEG, Biopac Systems Inc., n.d.) was placed in a sleeve and attached to participants’ right arm with a strap. The preparation procedure required 15 minutes prior to participant arrival and between 15 and 30 minutes in the presence of the participant (30-45 minutes in total).

The electrode cables were quite short, rested loosely at the participant’s back and were attached to the amplifier. The amplifier wirelessly transferred the EEG signal through a USB Wi-Fi antenna to a recording laptop. The stimulus computer was linked via its parallel port to the Digital I/O (37 pin) port of an STP100C module (isolated digital interface) attached to the MP160 Biopac device (Biopac Systems, Inc., n.d.) which allowed for the digital event markers to be recorded using Acqknowledge. However, the EEG data stream from the Mobita amplifier and the event marker data stream from MP160 could not be integrated into one recording pane or synchronised across two recording panes in Acknowledge (version 5.0.3). To solve this, a bespoke setup was made, whereby the event markers were sent via a wired connection to the Mobita amplifier to integrate into the recording at source. The integrated (EEG & event marker) data was then transferred wirelessly to the recording laptop as described before. Further details on this setup can be found in the supplementary file. Recordings were performed in an unshielded room as the system has been designed to be mobile and suitable for use in a wide range of environments. Table 1 displays a summary of technical differences between the two systems.

**Table 1.** A summary of technical differences between the EasyCap and Mobita EEG systems.

|  | EasyCap | Mobita |
| --- | --- | --- |
| **Non-Overlapping Channels** | TP9, TP10 | FPz, Oz |
| **Ground Electrode Location** | AFz | Middle of the forehead |
| **Reference Electrodes' Location** | Mastoid, external to the cap | Mastoid, within the cap |
| **Reference Recording Mode** | Linked Mastoids (Implicit) | Common Average Reference |
| **Electrooculographic Electrodes** | Separate HEOG and VEOG | Fp1, F7 |
| **Electrode Impedance** | <5kΩ | Not available |
| **Online Filter** | 250Hz | None |
| **Sampling Rate** | 500Hz | 1000Hz |
| **EEG Cap Size** | Based on the head size | One adjustable size |
| **Total Preparation Time** | 30-60 min | 30-45 min |
| **Recording Room** | Shielded | Unshielded |

#### 2.2.3. Cognitive task

The participants completed an arrow version of the flanker task (Eriksen and Eriksen, 1974) whilst the EEG data was acquired. This is a commonly used task in the study of attentional and error-control processes suitable for ERP research investigating both stimulus- and response-locked components such as P300 and error-related negativity (ERN; Pratt et al., 2011; Rietdijk et al., 2014). The task was presented using E-Prime software version 3 (Psychology Software Tools, 2012). Each trial consisted of 7 arrowheads presented at the centre of the screen. The target stimulus was the middle arrowhead and participant’s task was to detect whether it was pointing left or right and respond using the computer keyboard (letter “C” for left and letter “M” for right). Three distractor arrowheads on each side of the target changed direction depending on the trial condition. If they pointed in the same direction as the target the condition was congruent. If in the opposite direction, it was incongruent. In the neutral condition the distractor arrowheads were replaced with the letter “v”. Each trial was proceeded by a fixation cross. Maximum response time in each trial was 600ms and between-trial intervals were jittered in duration (400-1600ms). One participant who initially participated in the study during the pilot completed 750 trials and all remaining participants completed 600 trials (200 per condition) with the task taking approximately 20 minutes.

#### 2.2.4. Researcher and participant experience

Participants’ experiences of both systems were discussed at phase two. Their observations were noted retrospectively and remain anecdotal in nature. However, they provide important practical information which should be considered for research protocols involving the Mobita system. Researcher notes focused on technical issues observed during the recordings. All observation notes can be found in the Demographics file in the project repository (https://osf.io/kubv5/; Topor et al., 2021).

### 2.3. Data Analyses

#### 2.3.1. Data import and digital marker positions

Offline analysis of the EEG data from both systems was performed using BrainVision Analyzer 2 (Brain Products, 2012). EasyCap data were recorded in a format compatible with BrainVision. Digital event markers were integrated and correctly numbered to reflect different event types (congruent, incongruent, and neutral stimuli). Correct and incorrect responses were marked using participant response data extracted from Eprime and a Perl script that was previously prepared and used with the task.

EEG data recorded with Mobita were exported to.EDF format (Kemp et al., 1992) and imported to BrainVision Analyzer. As a result of the bespoke solution for digital signal recording, only values 0 and 1 could be registered to mark events. Therefore, all stimuli events were marked when a change from 0 to 1 occurred in the digital channel and all responses were marked when 1 changed back to 0. In R Studio (RStudio Team, 2020), task relevant data recorded with Eprime were used alongside the Acqknowledge markers to label the type of condition and correct/incorrect responses.

For a detailed description of the preparation of digital markers for the bespoke digital signal transfer used for Mobita in this study, see the supplementary material. One particularly significant difficulty observed during this process was data loss in the Mobita recordings of three participants caused by two types of signal drops. The first type was caused by Wi-Fi connection issues and led to no data being recorded (all channels were flat) for the duration of a few trials at each instance. The second type of signal drop led to a complete termination of recording in the Acknowledge software for about one minute in one case. The cause of this is unknown. Acqknowledge does not record the duration of recording termination in such situations, and this had to be determined manually by comparing the timings recorded by Acqknowledge and Eprime and determining the temporal location of the gap. Data loss in this case included 68.09 seconds of data (40 consecutive trials).

#### 2.3.2. Pre-processing

For pre-processing, only channels overlapping between the two systems were selected. In EasyCap, TP9 and TP10 and in Mobita, Fpz and Oz were excluded. Data were visually inspected for channels with no or extreme activity. No channels were interpolated for EasyCap. In Mobita, channels were interpolated in two recordings (one channel in the first case and two channels in the second case). In addition, during the inspection of the Mobita data, the mastoid reference channels were observed to be extremely noisy or flat in three recordings. Therefore, all EEG recordings from both systems were re-referenced to the average activity of the subset of overlapping channels (for EasyCap, this included the initial implicit reference). Subsequently, the following filters were applied: 0.1 Hz high-pass, 50Hz low-pass and 50Hz notch filter with threshold selection designed to avoid ERP distortion and ensure the most optimal signal to noise levels based on best practice recommendations and previous EEG system comparisons (Tanner et al., 2015, 2016). Data were then re-sampled to 512 Hz for both systems. Before artefact cleaning, all non-task data were removed. This included the start and the end of the recordings as well as breaks between the blocks leaving only task-related block segments for further analysis.

Ocular correction independent component analysis was used with default BrainVision Analyzer settings (Brain Products GmbH, 2019, p. 279) to automatically detect components around blinks. Channels used to train the algorithm were HEOG and VEOG in EasyCap and Fp1 and F7 in Mobita. Component rejection was semi-automatic where one researcher (MT) inspected each ICA component and confirmed its removal/retainment. There were no significant differences between the number of components removed in EasyCap (*Median* = 2.5, *IQR*= 1) and Mobita (*Median =* 2.5, *IQR=*3; *V*=7.5, *p*=.59). Data were epoched into two types including stimulus-locked epochs for frequency and P300 analyses and response-locked epochs with correct and incorrect responses for ERN analyses. Stimulus-locked epochs for frequency and P300 analyses were selected at −200ms to 500ms respective to stimulus onset. Response-locked epochs for the ERN analysis were selected at −150ms and 200ms respective to response onset. Automatic artefact rejection was performed on all epochs using the default settings of BrainVision Analyzer excluding trials with activity below 0.5μV for a duration of 50ms, amplitude values falling outside of the −200μV and 200μV range, absolute amplitude difference above 200μV in any interval of 200ms and lastly, voltage steps of more than 50μV per millisecond.

#### 2.3.3. Noise Measurements

To assess potential noise in raw data, the fast Fourier transform was applied to unfiltered data that were re-referenced, re-sampled and segmented to task-related blocks without any ocular correction or artefact rejection. 0.1-2Hz and 49-51Hz power values were then extracted for further analysis to understand the potential of slow drift interference (de Cheveigné and Arzounian, 2018) and line noise interference (Leske and Dalal, 2019).

Levels of noise were also assessed in the fully pre-processed and cleaned stimulus-locked data. First, the fast Fourier transform was applied and power values at 0.1-2Hz and 49-51Hz were extracted for the comparison of slow drift and line noise interference before and after pre-processing. In addition, the proportion of rejected artefactual stimulus-locked trials was calculated for both systems. Next, signal to noise ratio (SNR) and the root mean square (RMS) values were calculated from a subset of electrodes excluding those located at the edges of the cap which are particularly prone to noise. The remaining subset therefore included F3, Fz, F4, FC5, FC1, FC2, FC6, C3, Cz, C4, CP5, CP1, CP2, CP6, P3, Pz, P4. SNR and RMS metrics are common in studies comparing different types of EEG equipment (Kam et al., 2019; Mathewson et al., 2017; Oliveira et al., 2016). SNR was calculated from averaged, stimulus-locked trials using the formula embedded within the BrainVision Analyzer software (Brain Products GmbH, 2019, p. 402) which estimates the average signal power as squared absolute values of the average data across all data points and all frequency bins whilst noise power is estimated as the biased variance of the data across all segments. The values were extracted for all electrodes included in the specified subset and then averaged for each participant. RMS values were determined with BrainVision Analyzer’s RMS function which calculated the root from the average of the squares of the individual values (Brain Products GmbH, 2019, p. 210) within the stimulus-locked epochs’ baseline period of −200 to −100ms prior to stimulus onset.

#### 2.3.4. Time-Frequency Measurements

For time-frequency analyses of task-related brain activity, the fast Fourier transform was applied to the pre-processed stimulus-locked epochs. Power was extracted from the same subset of electrodes used in the SNR and RMS analyses (see previous section). The data were analysed in four frequency bands including theta (4-8Hz), alpha (8-14Hz), low Beta (14-24Hz) and high beta (24-30Hz). For the comparison of power activity within these bands, we followed the method used by Kam et al. (2019) whereby 5 electrodes with maximum activity were identified per system and power was averaged across the overlapping channels. For theta and alpha, electrodes with the most positive power were selected (theta Fz, F4, FC1, FC2, alpha Fz, Pz, P4) due to expected engagement of cognitive control and attentional processes. For low and high beta, electrodes with the least positive power were selected (low beta CP1, CP2, Pz, high beta CP1, CP2, Pz, P4) as pre-response motor-related beta desynchronisation was expected (Doyle et al., 2005).

#### 2.3.5. Event-Related Potential Measurements

For ERP analyses, baseline correction was applied to all fully pre-processed epochs. The baseline window was located at −200ms and −100ms prior to stimulus onset for stimulus-locked P300 epochs and at −150ms to −100ms for response-locked ERN epochs following best practice recommendations by Alday (2019). Subsequently, the number of included epochs for each type (stimulus-locked and response-locked) was matched for each participant across the two systems. Additionally, the number of response-locked epochs was matched between correct and incorrect responses. The selection of trials for matching was based on the order of occurrence. For one participant, there were only 5 trials with incorrect responses available for ERN analyses. This is below the recommended value of at least six trials (Olvet and Hajcak, 2009) so this case was removed from all ERN analyses. EEG activity was then averaged across trials. Mean amplitude for the P300 component was extracted for a 300 to 500ms interval at Pz and for the ERN component for a 0 to 100ms interval at Cz which is suitable for flanker task analyses (Klawohn et al., 2020; Rietdijk et al., 2014). To investigate the characteristics of the obtained ERPs, peak amplitude and latency were also extracted. ERN and P300 peaks were semi-automatically identified in BrainVision Analyzer.

#### 2.3.6. Statistical Analyses

All dependent variables were tested using the Wilcoxon test to assess central tendency differences and Fligner-Killeen test to assess homogeneity of variance between the two systems. The dependent variables were divided into the following groups for the purpose of controlling the familywise error: average power values at 0.1-2Hz and 49-51Hz before and after pre-processing; proportion of artefactual stimulus-locked trials; stimulus-locked noise level metrics including SNR and RMS; averaged power for four frequency bands (theta, alpha, low beta, high beta); mean amplitude, peak amplitude and peak latency for P300 and ERN. Bonferroni correction was applied accordingly.

In addition to tests of difference, averaged power for each frequency as well as the mean amplitude values for P300 and ERN were correlated between the two systems using Kendall’s Tau correlation. These analyses were also adjusted using the Bonferroni correction.

##### 2.3.6.1. Exploratory analyses

Due to the observed P300 and ERN topographical differences between the two systems, we decided to explore whether SNR values might differ between the two systems by electrode locations. We therefore divided the SNR electrode subset into three general regions: frontal (Fp1, F2p, F7, F3, Fz, F4, F8), central (FC5, FC1, FC2, FC6, C3, Cz, C4, CP5, CP1, CP2, CP6) and posterior (P7, P3, Pz, P4, P8, O1, O2). We calculated the average SNR values for these regions for each participant and each system and compared these values using the Wilcoxon and Fligner-Killeen tests. Bonferroni correction was applied.

##### 2.3.6.2. Power Analysis

The results of the study showed some significant differences between the EEG systems, but also a number of potentially practically informative non-significant differences with large effect sizes. This could be due to the small sample size, the large number of statistical tests and the Bonferroni correction for multiple comparisons. To help in the interpretation of these potentially informative effects, we decided to run a post-hoc power analysis on the smallest large effect size obtained. Post-hoc power analyses are discouraged when reliant on effect sizes achieved with a limited sample as it is not possible to estimate whether these effect sizes reflect the true population effects (Lakens, 2021). However, considering the methodological focus of the current study and the practical importance of differences between the two systems, it is important to understand the current study’s statistical sensitivity to detect the effects of interest. The power calculation was conducted with G*Power (Faul et al., 2007) using the results of the 0.1-2Hz raw data power comparison between EasyCap and Mobita where a non-significant large effect size was found (*r*=.53). The analysis yielded power of 21.7% (calculation output https://osf.io/fdmzx/). This therefore reflects very small chances for obtaining statistically significant results, even for large effect sizes of interest with the current sample and with the given number of tests and comparisons. To aid the interpretation of the results, we deemed all results with large effect sizes as practically informative regardless of whether the p-values reached the desired 0.05 threshold.

## 3. Results

### 3.1. Researcher and Participant Experience

For the fit of the cap, during the EasyCap recording, common comments referred to the chin strap that felt “scratchy” for some participants. It had to be adjusted throughout the procedure to improve comfort. During the Mobita recording, in some instances the front of the cap put pressure on the forehead which led to moderate discomfort. The Mobita cap has a tightening string which helps to adjust the fit though it is positioned around the face only. We either loosened it up for participants or refrained from using it to improve comfort.

Participants were generally impressed with Mobita due to its shorter preparation time. The preparation procedure that involved participants took up to 30 minutes compared with up to an hour for EasyCap. Some were also relieved that they did not have to wash their hair following the procedure and could quickly go back to their activities after participating in the study. One participant mentioned that they only agreed to participate again when they were informed that the procedure would be shorter this time and no gel would be used. It is worth noting however, that from the experimenter point of view, the total time taken to prepare each system was not very different as Mobita required extra preparation before participant arrival.

### 3.2. Noise Comparisons

Prior to pre-processing, average power was significantly more variable for Mobita at 0.1-2Hz (*X^2^*=8.9, *p*=.022) and 49-51Hz (*X^2^*=11.0, *p*=.007) compared to EasyCap but there were no significant differences for central tendency comparisons. However, there was a large effect size for the difference in central tendency at 0.1-2Hz (*r*=.53) indicating that Mobita may be more likely to yield larger power values. Figure 2 displays individual data plots with average power at 0.1-2Hz and 49-51Hz for both systems as well as the overall averaged power spectrum for the raw EEG data.

**Figure 2.**
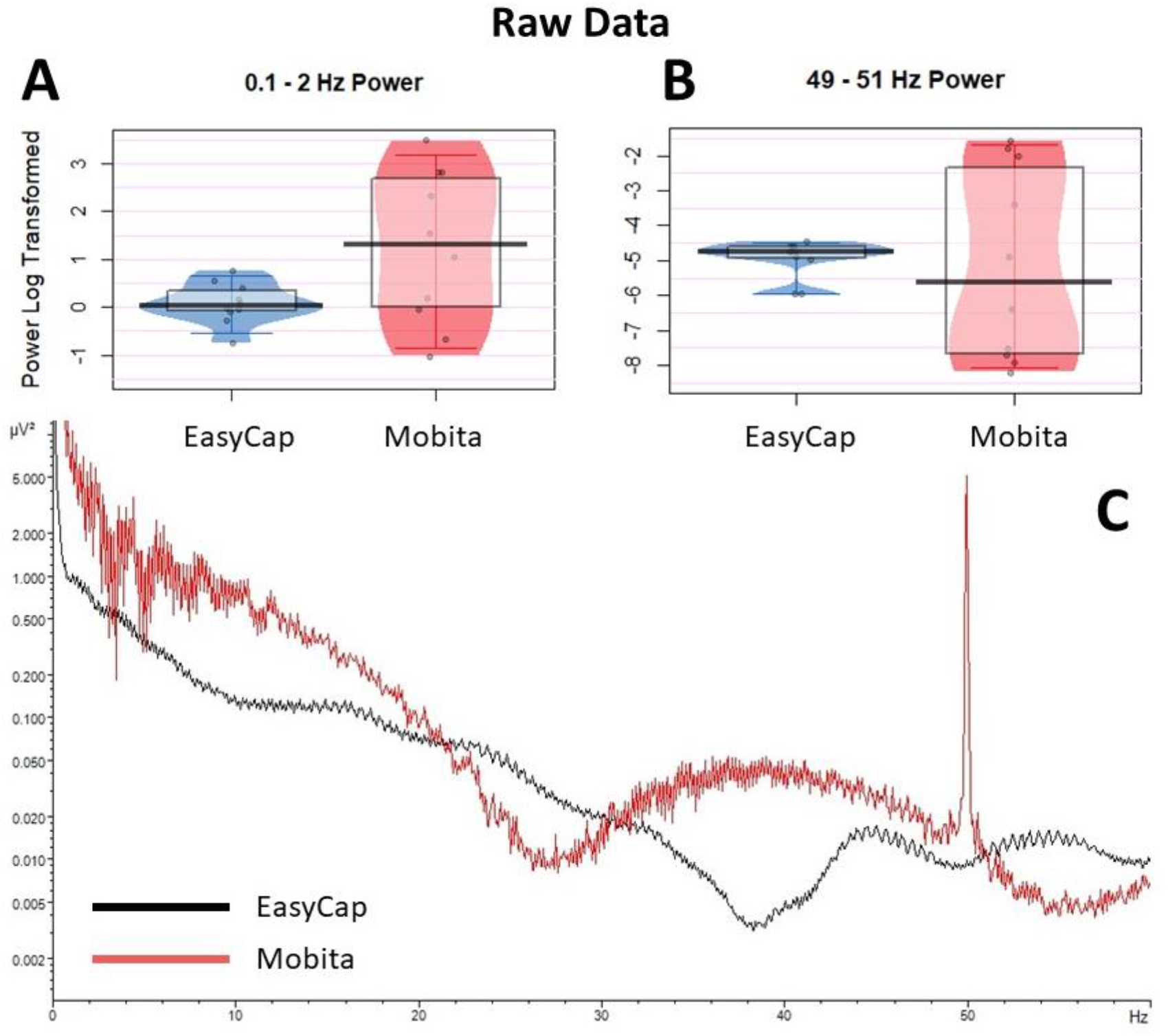
Raw data average power prior to pre-processing. Jittered individual data points are plotted for 0.1-2Hz **(A)** and 49-51Hz **(B)** to compare between the EasyCap and Mobita recordings. The vertical bar marks the median and the shaded box reflects the inter-quartile range. **C** Is a representation of log-transformed power spectrum at 0 to 60 Hz for EasyCap and Mobita.

Following pre-processing, there were no statistically significant differences in central tendency or variance between the two systems. However, at 0.1-2Hz, the Fligner-Killeen *X*^2^ value only decreased slightly and the difference in variance is nearing the significant p-value threshold. Figure 3 displays individual data plots with average power at 0.1-2Hz and 49-51Hz for both systems as well as the overall averaged power spectrum for the pre-processed EEG data. Exact statistical results are presented in Table 2.

**Figure 3.**
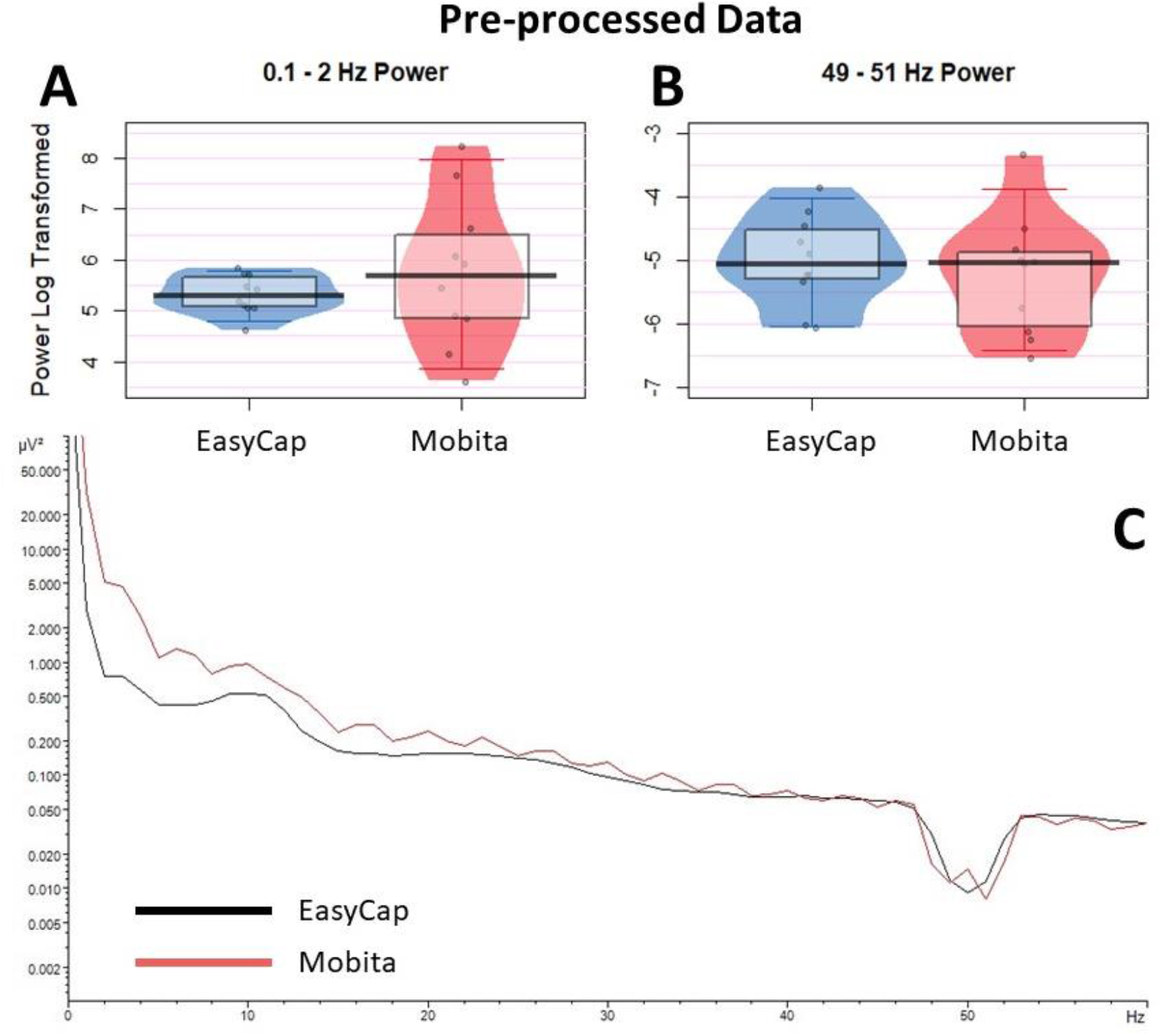
Pre-processed data average power. Jittered individual data points are plotted for 0.1-2Hz **(A)** and 49-51Hz **(B)** to compare between the EasyCap and Mobita recordings. The vertical bar marks the median and the shaded box reflects the inter-quartile range. **C** Is a representation of log-transformed power spectrum at 0 to 60 Hz for EasyCap and Mobita.

**Table 2.** Median and inter-quartile ranges are displayed for both systems before and after pre-processing for average power at 0.1-2Hz and 49-51Hz. The results of statistical comparisons of the central tendency (Wilcoxon) and variance (Fligner-Killeen) are also presented. Significant differences are marked with an asterisk. Bonferroni correction was used to adjust the obtained p-values.

| | EasyCap<br>Median (IQR) | Mobita<br>Median (IQR) | Central Tendency<br>Wilcoxon<br>$V, p (r)$ | Variance<br>Fligner-Killeen<br>$X^2, p$ |
| --- | --- | --- | --- | --- |
| **Raw Data** |  |  |  |  |
| **0.1-2Hz** | 0.03 (0.42) | 1.31 (2.67) | 11, .84 (.53) | 8.90, .022* |
| **49-51Hz** | -4.76 (0.33) | -5.64 (5.30) | 30, 1.0 (.08) | 11.01, .007* |
| **Pre-processed Data** |  |  |  |  |
| **0.1-2Hz** | 5.31 (0.57) | 5.68 (1.61) | 18, 1.0 (.31) | 6.86, .07 |
| **49-51Hz** | -5.06 (0.78) | -5.05(1.16) | 32, 1.0 (.21) | 0.44, 1.0 |

Artefact rejection rates for Mobita were significantly higher (*V*=1, *p*=.008, *r*=.85) and more variable (*X^2^*=6.42, *p*=.023) than for EasyCap. There were no statistically significant differences between the two systems in terms of SNR central tendency or variance. Baseline RMS was significantly more variable for Mobita compared with EasyCap (*X^2^*=12.74, *p*=.001) but there was no difference in terms of central tendency. However, all central tendency noise comparisons (artefact rejection, SNR and RMS) between the two systems yielded large effect sizes indicating a possibility that Mobita recordings may have lower SNR and higher baseline RMS in comparison to EasyCap. Exact statistical results are reported in Table 3 and individual data plots are presented in Figure 4.

**Table 3.**
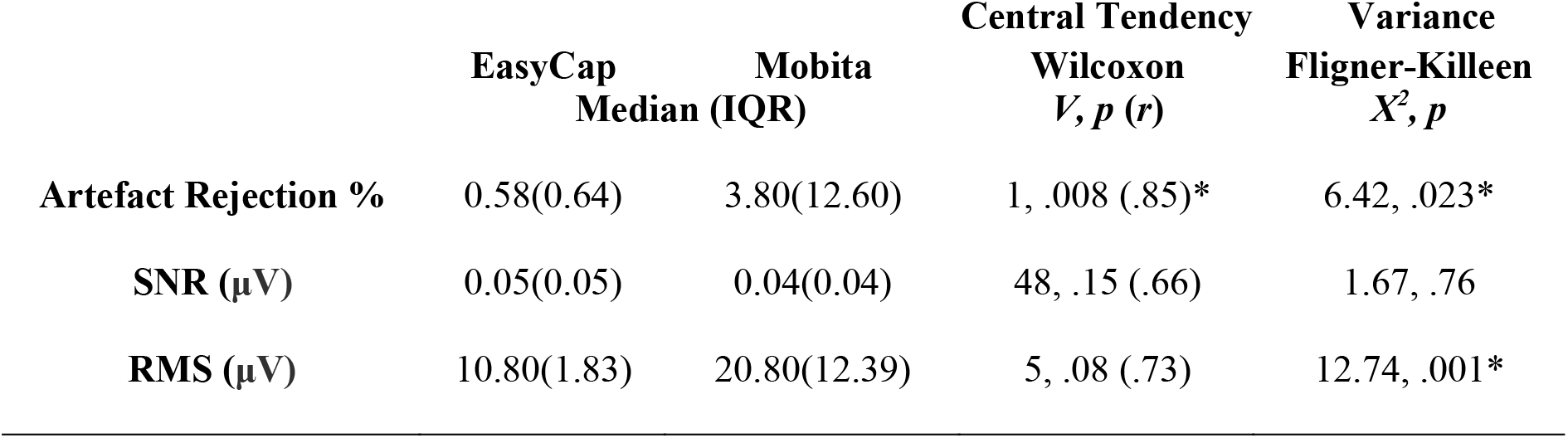
Median and inter-quartile ranges are displayed for both systems for all three measures of noise: percentage of rejected artefactual trials, signal-to-noise ratio, and root mean square. The results of statistical comparisons of the central tendency (Wilcoxon) and variance (Fligner-Killeen) are also presented. Significant differences are marked with an asterisk. Bonferroni correction was used to adjust obtained p-values.

| | EasyCap<br>Median (IQR) | Mobita<br>Median (IQR) | Central Tendency<br>Wilcoxon<br>$V, p (r)$ | Variance<br>Fligner-Killeen<br>$X^2, p$ |
| --- | --- | --- | --- | --- |
| **Artefact Rejection %** | 0.58(0.64) | 3.80(12.60) | 1, .008 (.85)* | 6.42, .023* |
| **SNR (μV)** | 0.05(0.05) | 0.04(0.04) | 48, .15 (.66) | 1.67, .76 |
| **RMS (μV)** | 10.80(1.83) | 20.80(12.39) | 5, .08 (.73) | 12.74, .001* |

**Figure 4.**
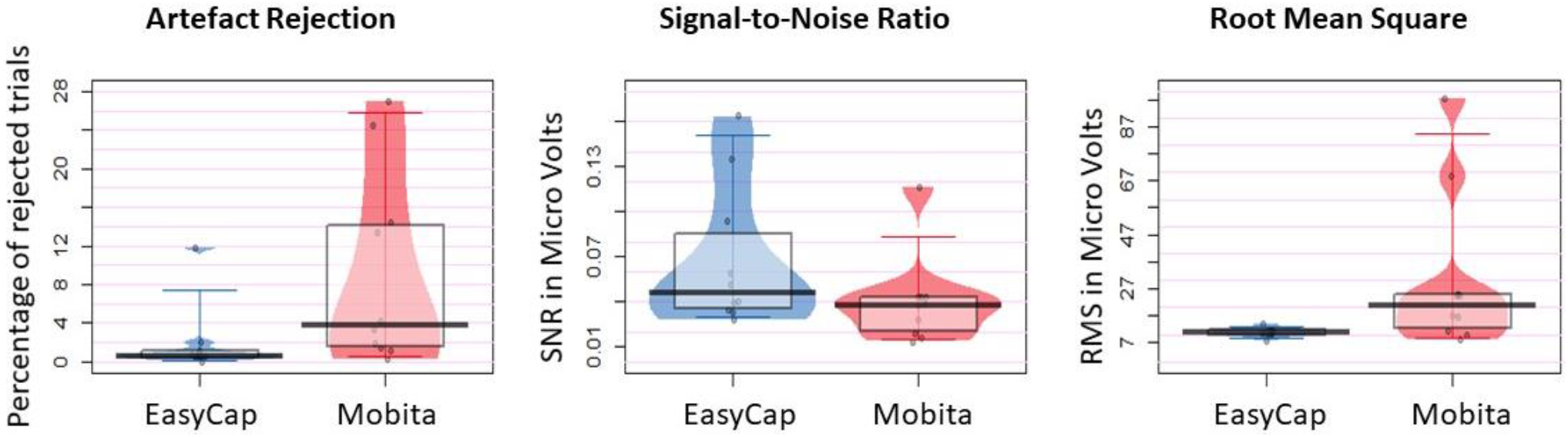
Jittered individual data points reflect average percentage of rejected artefactual trials, signal-to-noise ratio and root mean square for EasyCap and Mobita recordings. The vertical bar marks the median and the shaded box reflects the inter-quartile range.

### 3.3. Frequency Power Comparisons

Frequency power was compared between EasyCap and Mobita across four bands (theta, alpha, low beta and high beta). No statistically significant results were obtained for the tests of difference in central tendency and variance and the correlations were also non-significant. However, the central tendency differences between the two systems yielded large effect sizes for low and high beta power values where weaker activity has been recorded with Mobita. In addition, a large correlation was observed between EasyCap and Mobita for the high beta band. Medians, inter-quartile ranges and exact test results can be found in Table 4. Figure 5 displays topographical power distribution, scatter plots and individual data plots for EasyCap and Mobita across the four frequency bands.

**Table 4.** Median and inter-quartile ranges are displayed for both systems for four frequency bands: theta, alpha, low beta and high beta. The results of statistical comparisons of the central tendency (Wilcoxon), variance (Fligner-Killeen) and the correlations (Kendall’s Tau) between the two systems are also displayed. Bonferroni correction was used to adjust the obtained p-values.

|  | EasyCap<br>Median (IQR) | Mobita<br>Median (IQR) | Central Tendency<br>Wilcoxon<br>*V, p, (r)* | Variance<br>Fligner-Killeen<br>*X<sup>2</sup>, p* | Correlation<br>Kendall's Tau<br>*τ, p* |
| --- | --- | --- | --- | --- | --- |
| **Theta** | -0.78(0.59) | 0.10(1.03) | 12, 1.0, (.50) | 1.03, 1.0 | .11, 1.0 |
| **Alpha** | -1.15(0.88) | -0.38(1.82) | 18, 1.0, (.31) | 1.23, 1.0 | .42, 1.0 |
| **Low Beta** | -2.35(0.67) | -1.65(0.82) | 6, .33, (.69) | 1.72, 1.0 | .42, 1.0 |
| **High Beta** | -3.67(0.84) | -3.11(0.67) | 3, .12, (.79) | 0.07, 1.0 | .51, .56 |

**Figure 5.**
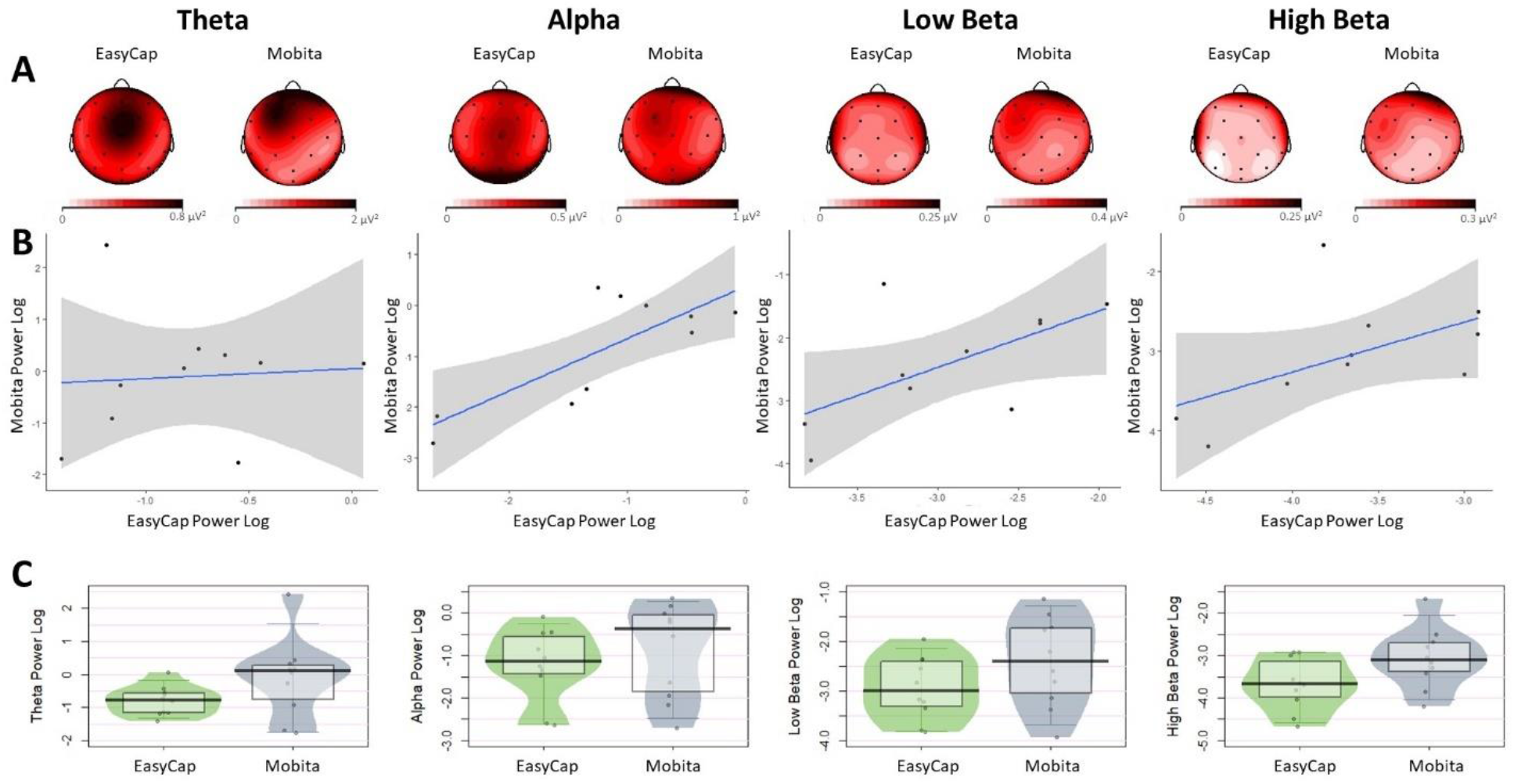
**A.** Topographies for all power frequencies are displayed for comparison between the two systems. The topographies have not been normalised and different scales are used for EasyCap and Mobita. **B.** Scatter plots with fitted line of best fit and confidence intervals to visually reflect the relationship between power obtained with the EasyCap and Mobita systems. **C.** Jittered individual data points reflecting average power for each participant recorded with each system. The vertical bar marks the median and the shaded box reflects the inter-quartile range.

### 3.4. Event-Related Potentials Comparisons

#### 3.4.1. P300

No statistically significant differences were found between EasyCap and Mobita for the mean P300 amplitude at 300ms to 500ms in terms of central tendency and variance. All observed effect sizes were small. The correlation between the two systems was also non-significant with a small-medium relationship. In addition, no statistically significant differences in central tendency or variance were identified in P300 peak amplitude or peak latency. Table 5 displays the medians and interquartile ranges observed, as well as exact statistical results. Figure 6 displays the P300 waveforms, topographies, a scatter plot and an individual data plot for comparison of P300 mean amplitude values.

**Table 5.** Median and inter-quartile ranges are displayed for both systems for the measures of P300 mean amplitude at 300ms to 500ms, ERN mean amplitude at 0ms to 100ms, peak amplitude values and peak latency. The results of statistical comparisons of the central tendency (Wilcoxon) and variance (Fligner-Killeen) are displayed for all measures. In addition, the mean amplitude correlation results (Kendall’s Tau) are displayed. Bonferroni correction was used to adjust all obtained p-values.

|  | EasyCap<br>Median (IQR) | Mobita | Central Tendency<br>Wilcoxon<br>*V, p, r* | Variance<br>Fligner-Killeen<br>*X<sup>2</sup>, p* | Correlation<br>Kendall's Tau<br>*τ, p* |
| --- | --- | --- | --- | --- | --- |
| **P300** |  |  |  |  |  |
| Mean Amplitude (μV) | 2.40 (1.63) | 1.69 (2.39) | 31, 1.0 (.11) | 1.53, 1.0 | 0.24, 1.0 |
| Peak Amplitude (μV) | 2.54 (1.75) | 2.49 (3.22) | 21, 1.0 (.15) | 1.69, 1.0 |  |
| Peak Latency (ms) | 346 (38) | 343 (67) | 23, 1.0 (.16) | 2.85, 1.0 |  |
| **ERN** |  |  |  |  |  |
| Mean Amplitude (μV) | -2.69 (5.46) | -0.89 (1.62) | 1, .11 (.85) | 1.04, 1.0 | .28, 1.0 |
| Peak Amplitude (μV) | -3.96 (7.96) | -4.56 (0.98) | 36, 1.0 (.02) | 0.31, 1.0 |  |
| Peak Latency (s) | 41 (16) | 6 (29) | 13, .20 (.87) | 1.48, 1.0 |  |

**Figure 6.**
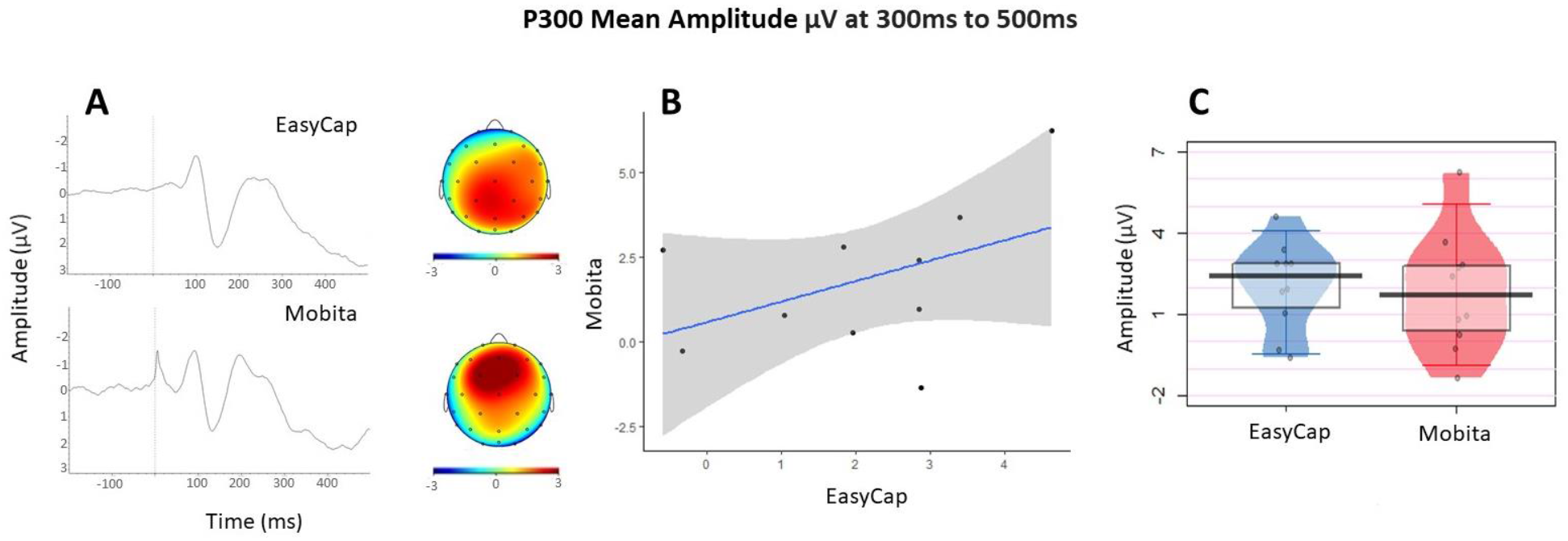
**A.** The P300 waveforms and topographies for each system. **B.** Mean amplitude scatter plot with a fitted line of best fit and confidence intervals to visually reflect the relationship between EasyCap and Mobita. **C.** Jittered individual data points reflecting average mean amplitude for each participant recorded with each system. The vertical bar marks the median and the shaded box reflects the inter-quartile range.

#### 3.4.2. ERN

No statistically significant differences were found between EasyCap and Mobita for the mean ERN amplitude at 0ms to 100ms in terms of central tendency and variance. The correlation between the two systems was also non-significant with a small-medium association. In addition, no statistically significant differences in central tendency or variance were identified in ERN peak amplitude or peak latency. However, mean amplitude and peak latency central tendency differences between the two systems yielded large effect sizes. In Figure 6A it is evident that the ERN peak occurs early in Mobita, almost directly at the time of response onset. Table 5 displays the medians and interquartile ranges observed, as well as exact statistical results. Figure 6 displays the P300 waveforms, topographies, a scatter plot and an individual data plot for comparison of ERN mean amplitude values.

### 3.5. Exploratory Analysis

Exploratory analysis focused on the differences in SNR values by electrode locations including frontal, central and posterior. In Figures 6A and 7A, we have observed that the obtained ERP peaks are shifted frontally. The shift is especially prominent for P300 (Figure 6A). We were therefore interested in finding out whether some electrodes might be particularly susceptible to high noise levels. We found no differences in variance between the two systems. However, all central tendency differences between EasyCap and Mobita yielded large effect sizes with this difference being statistically significant for posterior electrodes (*V*=54, *p*=.04, *r*=.85). Table 6 displays the medians and interquartile ranges observed, as well as exact statistical results. Figure 7 presents the individual data plot for comparison of SNR values across electrode location and between EasyCap and Mobita.

**Figure 7.**
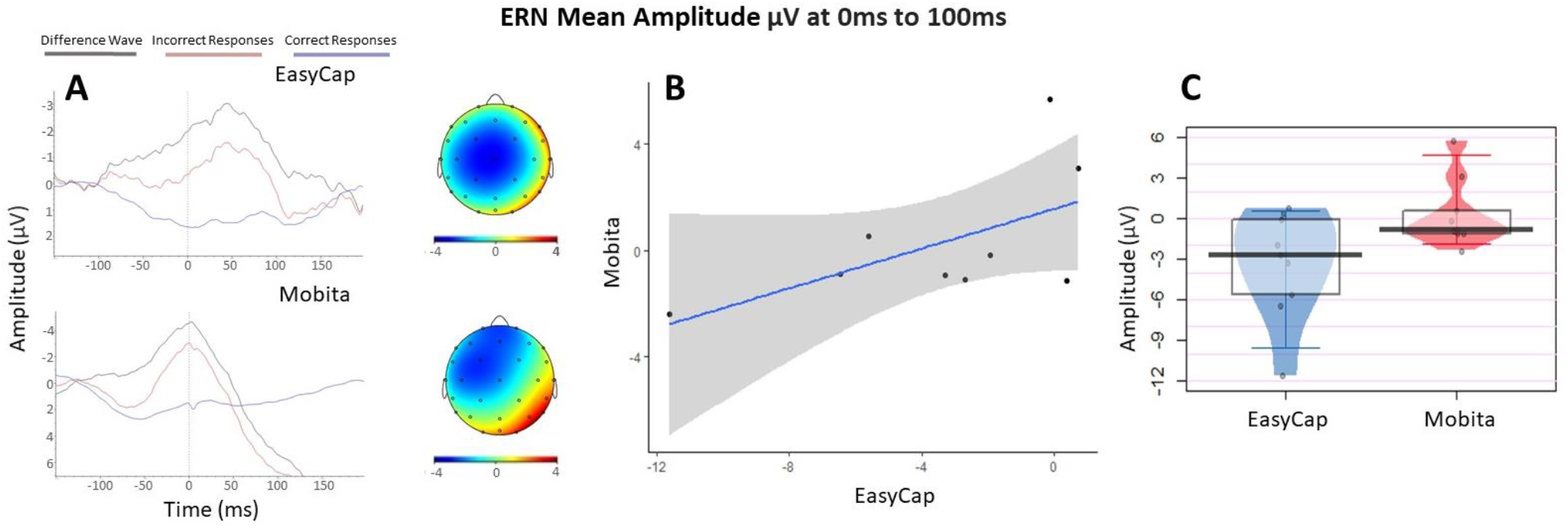
**A.** The ERN waveforms and topographies for each system. **B.** Mean amplitude scatter plot with a fitted line of best fit and confidence intervals to visually reflect the relationship between EasyCap and Mobita. **C.** Jittered individual data points reflecting average mean amplitude for each participant recorded with each system. The vertical bar marks the median and the shaded box reflects the inter-quartile range.

**Table 6.** Median and inter-quartile ranges are displayed for both systems for SNR values recorded with frontal, central and posterior electrodes. The results of statistical comparisons of the central tendency (Wilcoxon) and variance (Fligner-Killeen) are provided. Statistically significant results are marked with an asterisk. Bonferroni correction was used to adjust all obtained p-values.

| SNR(μV) | EasyCap<br>Median (IQR) | Mobita | Central Tendency<br>Wilcoxon<br>$V, p (r)$ | Variance<br>Fligner-Killeen<br>$X^2, p$ |
| --- | --- | --- | --- | --- |
| **Frontal Electrodes** | 0.05(0.05) | 0.03(0.02) | 47, .49 (.63) | 0.46, 1.0 |
| **Central Electrodes** | 0.04(0.05) | 0.03(0.02) | 46, .63 (.60) | 0.33, 1.0 |
| **Posterior Electrodes** | 0.07(0.04) | 0.03(0.03) | 54, .04 (.85)* | 0.09, 1.0 |

## 4. Discussion

Mobita is an example of an innovative approach to EEG recordings via water-based electrodes and a wireless setup. These relatively new systems may be of interest to researchers who want to shorten the EEG preparation time (e.g., for children or clinical populations), reduce the possibility of sensory discomfort (e.g. for participants with sensory sensitivities) or considering independent recordings taken by participants or patients at home. From our experience, Mobita is currently not suited for a quick and easy application in studies aiming to analyse EEG recordings in time-locked epochs for frequency or ERP comparisons. Researchers considering the use of such systems should weigh the potential benefits against technical, practical and data quality disadvantages presented in this study.

### 4.1. Technical and Practical Considerations

Participants in the current study had a generally positive experience when using Mobita and some expressed their preference for Mobita over EasyCap due to the reduced preparation time and not having to wash their hair. However, none of the participants had any pre-existing sensory sensitivities, hyperactivity, or attention difficulties. It is not clear if Mobita would be more beneficial for participants with such difficulties and how much improvement it could bring overall to the experience of the EEG procedure.

From the researcher point of view however, the EasyCap system was more optimal in terms of the technical and practical application whilst the Mobita system required more adaptations and time-consuming solutions at all stages – set-up, recording and analysis. At set-up, it required a bespoke solution to allow for it to record synchronised digital and EEG signals for time-locked analyses. This is despite the setup being marketed as being able to record ERPs with no modifications or solutions that were alternative to the original expectations. The initial process of setting up Mobita is not straightforward and may require technicians or engineer assistance. At recording, the Mobita cap was not well fitted for some participants as one adjustable size was used instead of using caps fitted for individual head sizes like in the case of EasyCap. It was also problematic that the mastoid reference electrodes were embedded within the cap as it was difficult to keep them close to the scalp to obtain good quality reference data. As a result, an average reference was used during pre-processing instead. In addition, the unavailability of an electrode impedance measure meant that the researcher could not easily check and compare signal quality across the cap. Therefore, this increased the chances for Mobita EEG recordings to register more noise than EasyCap. Mobita was also susceptible to signal drop and recording termination which led to loss of data. During the analysis, the digital marker signal for cognitive task events (stimuli and responses) had to be extracted from Acqknowledge and labelled (condition and response types) externally whilst during the EasyCap recording the digital signal was mostly already labelled into different types. This was especially challenging for instances when the EEG and digital signal drops occurred as the gaps had to be manually detected and the markers were then realigned. Though this could be mitigated by choosing an option to record the signal directly to the logger instead of transferring the data wirelessly to the computer for recording. See the recommendation section below.

### 4.2. Data noise

Mobita had higher variance in registered noise at 0.1-2Hz and 49-51Hz in raw data compared to EasyCap. No other statistically significant differences were seen in raw or pre-processed data. A large effect size was observed in central tendency comparison of power at 0.1-2Hz in raw data between the two systems which indicates larger low-frequency drifts in Mobita compared to EasyCap. Following pre-processing, power at 49-51Hz became visually comparable with EasyCap (Figure 2). However, this was not the case for power at 0.1-2Hz, and power variance only reduced slightly numerically, with the statistical comparison remaining close to significance (Table 2, Figure 3A). This indicates the possibility that EEG data recorded with Mobita may be disadvantaged by slow drifts even after pre-processing. These drifts may be caused by poor electrode-to-skin contact and may mask slow cortical activity in studies looking at low frequencies or distort ERP components (de Cheveigné and Arzounian, 2018). In addition, we observed statistically significant central tendency and variance differences in the artefact rejection rates between the systems indicating higher noise levels and further data loss following the pre-processing procedures. SNR was generally lower for Mobita with large effect sizes obtained in comparisons of frontal, central and parietal electrodes and the latter reaching statistical significance. The RMS values were significantly more variable for Mobita and there was a large effect size observed for the central tendency comparison although it did not reach statistical significance. Taken together, these findings indicate that Mobita registers more noise at recording which can be to some extent improved with pre-processing. However, the high artifact rejection rates, low SNR and variable RMS suggest that the data will likely still contain higher levels of noise than standard EEG systems which may impact the EEG results as explained below.

### 4.3. EEG Results

Regarding the frequency analyses, power across the four bands (theta, alpha, low beta and high beta) did not significantly differ between the two systems. Power across the bands also did not significantly correlate between the systems. However, there was a large effect size detected for the difference in power between Mobita and EasyCap for the low and high beta frequency bands (Table 4, Figure 5). For these frequencies, we expected to observe motor-related beta desynchronization in the parietal regions reflected with power values that are negative or close to 0. Mobita activity seemed to be more positive than EasyCap activity potentially masking the motor-related beta desynchronisation. This might have been caused by significantly lower SNR in the parietal channels in the Mobita system as evident from the exploratory analysis (Figure 8). In addition, moderate to large τ correlations of power values were detected between the two systems for alpha and beta frequencies whilst there was no clear pattern of association for theta and the τ correlation was very small. This could be due to the observed increase in slow drift noise in Mobita (Figure 3). The topographies for all bands are relatively similar based on visual comparisons with a slight frontal shift of maximal power observed in theta and alpha bands recorded with Mobita.

**Figure 8.**
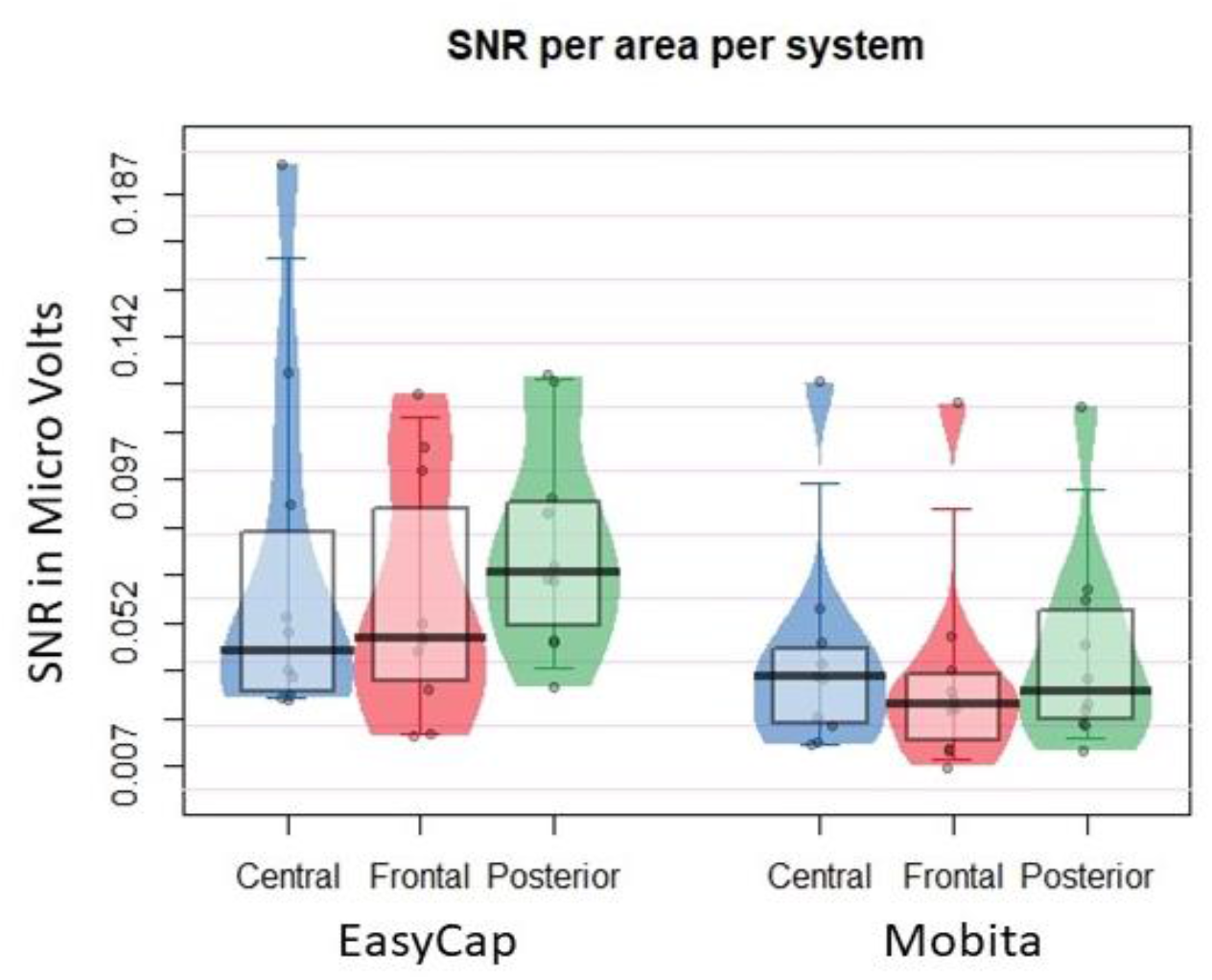
Jittered individual data points reflecting average SNR values for each participant presented by electrode location and recording system. The vertical bar marks the median and the shaded box reflects the inter-quartile range.

With regards to the ERP analyses, statistical comparisons between the two systems were non-significant with small effect sizes for P300 mean amplitude, peak amplitude and peak latency. The correlation of mean P300 amplitude values between the systems was small-medium τ and did not reach statistical significance. Therefore, there is no indication from the current data that Mobita could yield unreliable P300 results. The P300 waveforms seemed visually similar. However, looking at the P300 topographies (Figure 6B), EasyCap recorded maximal P300 activity over parietal regions which is expected in a Flanker task (Klawohn et al., 2020). In contrast, the P300 activity was maximal over the frontal region in Mobita which is unexpected. The topographies should be visually similar as the same participants were tested on the same task with both systems. As in the case of low and high beta power, it is possible that the posterior activity was masked by low SNR in parietal and centro-parietal channels which could have caused the shift of maximal P300 activity more frontally in Mobita. This is a significant issue for consideration in future research as it may lead to false interpretation of results which may be inconsistent with previous literature and the current understanding of frontal and parietal P300 variants (van Dinteren et al., 2014).

For ERN mean amplitude, peak amplitude and peak latency, there were no statistically significant differences between the two systems. The τ correlation of the ERN mean amplitude between the systems was small-medium and did not reach statistical significance. However, a large effect size was detected for the difference in central tendency mean amplitude at 0-100ms between the two systems with Mobita yielding lower amplitude values compared to EasyCap. Correspondingly, there was a large effect size for the peak latency central tendency comparison between the two systems which is also evident by visually inspecting the ERN waveform (Figure 7A). The Mobita ERN peak occurs almost at the onset of response, but it is normally expected at around 50ms post response onset in Flanker tasks (Klawohn et al., 2020; Riesel et al., 2013) which is accurately reflected in the EasyCap waveform. The likely explanation for this latency shift is the bespoke solution for digital marker recording used in Mobita. It seems that stimulus-locked events can be accurately marked with the current system where the digital signal is set from 0 to 1 at stimulus onset since the peak latency for the P300 component did not differ between the two systems. At response onset, the digital signal is set back from 1 to 0 and it seems that this event was recorded approximately 35ms earlier than the actual response if we directly compare the EasyCap and Mobita peak latency median values (Table 5). This issue may again have adverse consequences in future research and lead to false interpretation of results that will not be consistent with the current knowledge about the ERN. Otherwise, visual inspection of the ERN waveforms suggests that the ERN component registered with both systems is rather similar and occurs to be slightly clearer for Mobita than EasyCap. This may be due to practice effects as all Mobita recordings were taken during phase two of the study. The topographies are also visually similar with slight frontal shift observed in Mobita but this is unlikely to lead to false interpretation.

### 4.4. Limitations

One limitation of the current study is the small sample size and low statistical power. However, considering the practical nature of the study, even results that are statistically non-significant but have large effect sizes can be informative. The observed patterns and differences between the two systems will help researchers to understand the expected data quality obtained with water-based electrodes to avoid misinterpretation of results and to develop best practice solutions for those who decide to use such systems in the future.

Another limitation of the study is that in case of Mobita, it was the first time that the system was used to collect and analyse EEG data at the School of Psychology, University of Surrey. The EasyCap system has been used numerous times and best practices have already been established over the years. It is therefore possible that data recorded with EasyCap was of better quality because the researchers were able to use it more confidently. It is possible that with the development of best-practice solutions as well as further practice in the use of water-based systems, the recorded data will also be of higher quality. The current study is an important step in the development of these best practice solutions.

Lastly, authors have not conducted any formal recordings of the objective experiences at data acquisition for participants nor themselves. The experiences were retrieved from lab notes and memory. However, all described experiences illustrate difficulties which have led to certain practical adaptations or methodological considerations for the future as detailed in the section below. Therefore, the subjective nature of these experiences presents a case-scenario of a real-life application of a water-based EEG system.

### 4.5. Recommendations for future research

Based on the results presented in this study, we provide a set of general recommendations to avoid technical difficulties and false conclusions in studies using water-based electrode EEG systems. For more specific explanations regarding the assessment of electrode noise prior to recording, the bespoke solution on digital event markers and event markers labelling in the current study, see the supplementary file.

For studies using water-based EEG systems for low-frequency effects and ERPs, researchers may want to consider robust solutions for detrending data in order to prevent the slow drifts from distorting or masking the effects of interest. For studies focused on effects located in parietal regions, such as the P300 or beta desynchronisation, researchers should be especially careful about ensuring low noise levels. They should plan for regular assessments of data quality and perhaps consider taking more than one recording during a single procedure. The power spectrum of all channels can be assessed for noise during the recording breaks and the electrode fitting can be re-adjusted to improve the electrode to scalp contact. This would be especially beneficial for long procedures to ensure that the electrodes have not become dry or dislocated.

For studies requiring the use of mastoid reference channels, researchers should consider the likelihood of high noise levels due to the difficulty in sustaining good electrode to skin contact. One solution is to apply a bandage or an elastic band to secure the electrodes in place (see the supplementary file for an example).

Researcher should take a careful approach if their studies require good temporal precision (e.g., in ERP analyses) and if they are using a new EEG system which may or may not have been designed for time-locked analyses. To avoid possible latency shifts (as observed in this study), one solution is to synchronise the EEG recording with the stimulus presentation software at the point of the first stimulus onset and use event timing values from the stimulus recording software output rather than the digital signal.

Lastly, to avoid data loss due to signal drops in wireless systems, we recommend that researchers record EEG signal directly to the data logger instead of sending the data wirelessly to a recording computer. Issues associated with data loss and high noise levels can be further alleviated by aiming to recruit larger samples and increasing the number of trials in tasks to preserve statistical power.

### 4.6. Conclusions

Water-based EEG systems could potentially help to reduce participant involvement time and discomfort. However, they may require a number of time-consuming adaptations which are not necessary when using the state-of-the-art gel-based systems. Researchers should be aware of the likelihood that water-based systems will register high levels of noise which may impact analyses investigating ERPs, low frequencies and parietal activity. Otherwise, they may be at risk of drawing wrong conclusions from their results.

## 5. Author Contributions

The following contributions are specified following the CRediT taxonomy.

MT: conceptualisation, data curation, formal analysis, investigation, methodology, project administration, visualisation, writing – original draft, writing – review and editing.

BO: conceptualisation, supervision, verification, writing – review and editing.

PD: conceptualisation, analysis, methodology, supervision, verification, writing – review and editing.

All authors contributed to the article and approved the submitted version.

## 7. Acknowledgments

We would like to express our gratitude to all participants who have agreed to take part in this study; many thanks to student Paula Kreimeier who assisted with the management and analysis of the data at the early stages of the project and Ines Violante for providing invaluable support in the setup of the Mobita system as well as feedback on our plans for this manuscript.

## 8. Data Availability

All data, code, analysis outputs and supplementary materials for this project are deposited in an open-access repository which can be found at (https://osf.io/kubv5/; Topor, Opitz, & Dean 2021).

